# Developmental onset of enduring long-term potentiation in mouse hippocampus

**DOI:** 10.1101/787192

**Authors:** Olga I. Ostrovskaya, Guan Cao, Cagla Eroglu, Kristen M. Harris

## Abstract

Analysis of long-term potentiation (LTP) provides a powerful window into cellular mechanisms of learning and memory. Prior work shows late LTP (L-LTP), lasting >3 hours, occurs abruptly at postnatal day 12 (P12) in rat hippocampus. The goal here was to determine the developmental profile of synaptic plasticity leading to L-LTP in the mouse hippocampus. Two mouse strains and two mutations known to affect synaptic plasticity were chosen: C57BL/6 and *Fmr1*^*−/y*^ on the C57BL/6 background, and 129SVE and *Hevin*^−/−^ (*Sparcl1*^−/−^) on the 129SVE background. Like rats, hippocampal slices from all of the mice showed test pulse-induced depression early during development that was gradually resolved with maturation by 5 weeks. All the mouse strains showed a gradual progression between P10-P35 in the expression of short-term potentiation (STP), lasting ≤ one hour. In the 129SVE mice, L-LTP onset (>25% of slices) occurred by 3 weeks, reliable L-LTP (>50% slices) was achieved by 4 weeks, and *Hevin*^−/−^ advanced this profile by one week. In the C57BL/6 mice, L-LTP onset occurred significantly later, over 3-4 weeks, and reliability was not achieved until 5 weeks. Although some of the *Fmr1*^*−/y*^ mice showed L-LTP before 3 weeks, reliable L-LTP also was not achieved until 5 weeks. Two bouts of TBS separated by ≥90 minutes advanced the onset age of L-LTP in rats from P12 to P10. In contrast, L-LTP onset was not advanced in any of the mouse genotypes by multiple bouts of TBS at 90 or 180 minute intervals. These findings show important species differences in the onset of STP and L-LTP, which occur at the same age in rats but are sequentially acquired in mice.

**SIGNIFICANCE STATEMENT:** Long-term potentiation (LTP) is a cellular mechanism of learning and memory. Knowing the developmental profile for LTP provides a basis for investigating developmental abnormalities leading to intellectual disabilities and other neurodevelopmental disorders. Here we explore the developmental profile of LTP onset in two wild type mouse strains, C57BL/6 and 129SVE, together with *Fmr1*^*−/y*^ and *Hevin*^−/−^ (*Sparcl1*^−/−^) mutations that produce abnormalities in synaptic structure, plasticity, and development. Our data provide a foundation for future investigations into connections between structural and functional plasticity leading to developmental anomalies in the brain.

## INTRODUCTION

The hippocampus is critical for spatial navigation and processing of new information. It is the main brain region used to study long-term potentiation (LTP), a cellular mechanism of learning and memory. Knowing the maturational profile of synaptic plasticity provides a basis for investigating abnormalities leading to intellectual disabilities and other neurodevelopmental disorders. Our previous work on Long-Evans rats revealed test pulse depression that lasts until postnatal day 21 (P21) (Cao & Harris, 2012), replicating earlier findings (Abrahamsson, Gustafsson, & Hanse, 2007, 2008). Theta-burst stimulation (TBS) reversed the test pulse depression at P8 to P11, but no potentiation was produced above the initial naïve response. At P12, the TBS reliably induced enduring LTP lasting more than 3 hours (L-LTP). When multiple episodes of TBS were delivered, the onset age of L-LTP was advanced to P10 (Cao & Harris, 2012). Here our goal was to extend these studies to mouse hippocampus.

Mice are a widely used model system to test the effects of genetic manipulations on normal behavior and physiology (Ellenbroek & Youn, 2016; Homberg, Wohr, & Alenina, 2017). Little is known about the developmental profile of synaptic plasticity in mice. Two commonly used wild type mouse strains, C57BL/6 and 129SVE, were chosen together with *Fmr1*^*−/y*^ and *Hevin*^−/−^ (*Sparcl1*^−/−^), which are known for producing aberrations in synaptic plasticity and development. *Fmr1*^*−/y*^ on the C57BL/6 background is a common model of Fragile X syndrome (FXS) for mental retardation and autism (He & Portera-Cailliau, 2013; Pfeiffer & Huber, 2009). The cause of FXS is the mutation preventing the synthesis of FMRP, an RNA-binding protein selectively expressed in neurons and responsible for mRNA transport and local protein synthesis in dendrites (Wang et al., 2016). Hevin is a protein released by astrocytes and interneurons that is critical for synapse formation and rearrangement (Mongredien et al., 2019). The synaptogenic activity of Hevin promotes glutamatergic synapse maturation and refines cortical connectivity and plasticity (Risher et al., 2014; Singh et al., 2016).

We tested for the impact of these two strains and two key mutations on the developmental profile of synaptic plasticity in hippocampal area CA1. The outcomes provide a foundation for investigating genetic effects on synaptic plasticity and may help to explain the inter-species variance in synaptogenesis and developmental capacity for learning and memory.

## MATERIALS AND METHODS

### Ethical Approval

Procedures were approved by the University of Texas at Austin Institutional Animal Care and Use Committee and complied with all NIH requirements for the humane care and use of laboratory mice (protocol # AUP-2012-00127, AUP-2012-00056, and their successor protocols).

### Animals

Breeding pairs of C57BL/6 (RRID:IMSR_JAX:000664) and *Fmr1*^*−/y*^ (RRID:MGI:5703659) on this background were kindly donated by Dr. D. Brager (Center for Learning and Memory, University of Texas at Austin) who received the founder pair from Dr. K. Huber (University of Texas Southwestern). Breeding pairs of 129SVES6 (129SVE) mice were obtained from a supplier (Taconic, Rensselaer, NY; RRID:IMSR_TAC:129sve), and *Hevin*^−/−^ (*Sparcl1*^−/−^, allelic composition *Sparcl1*^*tm1Pmc*^/*Sparcl1*^*tm1Pmc*^, RRID:MGI:4454665) were on this background. We will be referring to this knock-out as *Hevin*^−/−^. The generation of *Fmr1*^*−/y*^ and *Hevin*^−/−^ (*Sparcl1*^−/−^) has been described before (Barker et al., 2005; Consortium, 1994; McKinnon, McLaughlin, Kapsetaki, & Margolskee, 2000). Animals were co-housed and provided with food and water *ad libitum* on a 12 hr light-dark cycle. The experimental design was originally optimized for the *Fmr1*^*−/y*^ mice, in which the males have the strongest phenotypes; hence, for the appropriate comparison with *Fmr1*^*−/y*^ data, males were used for all the experiments. The exact age of each animal was known. Since the developmental profiles were more gradual in the mice than in rats, and for ease of graphical presentation, data from the mice were grouped by age as P10-13 (<2 wks), P13-17 (2 wks), P18-23 (3 wks), P26-31 (4 wks), and P32-37 (5 wks).

### Slice preparation

Hippocampal slices were prepared from mouse pups at P8 to P38 as previously described (J. N. Bourne, Kirov, Sorra, & Harris, 2007). Animals were decapitated under isoflurane anesthesia when appropriate (age older than P33). The brain was removed, and the left hippocampus was dissected out and rinsed with room temperature artificial cerebrospinal fluid (aCSF) containing (in mM) 117 NaCl, 5.3 KCl, 26 NaHCO_3_, 1 NaH_2_PO_4_, 2.5 CaCl_2_, 1.3 MgSO_4_, and 10 glucose, pH 7.4, and bubbled with 95% O_2_ / 5% CO_2_. Four slices (400 μm thick) from the middle third of the hippocampus were cut at 70° transverse to the long axis on a tissue chopper (Stoelting, Wood Dale, IL) and transferred to four individual interface chambers in the Synchroslice system (Lohmann Research Equipment, Castrop-Rauxel, Germany). The slices were placed on a net at the liquid-gas interface between aCSF and humidified 95% O_2_ / 5% CO_2_ atmosphere held at 32-33°C. The entire dissection and slice preparation took 5-7 min. The slices recovered in the chambers for 3 hr before the recordings commenced.

### Electrophysiology

The stimulation and data acquisition were obtained using the SynchroBrain software (Lohmann Research Equipment). A concentric bipolar stimulating electrode (FHC Inc., Bowdoin, ME) was positioned near the CA3 side, and a metal recording electrode (Thomas Recording, Geissen, Germany) was placed ~400 μm away from the stimulating electrode, also in the middle of CA1 *stratum radiatum*. Stimuli consisted of 200 μs biphasic current. An I/O curve was generated by measuring the slope (mV/ms) of the extracellular field excitatory postsynaptic potentials (fEPSPs) in response to increasing stimulus intensities (ranging from 100-500 μA). The I/O curve was used to determine the 50% response that was used and held constant for subsequent stimulation. The naïve fEPSP was the first response obtained at that 50% level several minutes after the completion of the I/O curve. The fEPSP slopes (mV/ms) were estimated by linear regression over the 0.2-0.4 ms interval in the middle of the initial negative slope of the fEPSP. This analysis time frame was kept constant for each slice throughout the recording. The stimulus intensity required to obtain the half maximal fEPSP slope was held constant for the duration of each experiment and applied every 5 min.

The various TBS paradigms are described in the figures and Results section. Briefly, the 8T TBS paradigm consisted of eight trains with 30 s intervals with each train containing 10 bursts at 5 Hz and each burst containing 4 pulses at 100 Hz. The 1T TBS paradigm consisted of one train of the same stimulation pattern. The fEPSP slope is expressed as a percentage of the naïve fEPSP or the averaged baseline response obtained 30 min before delivering the TBS paradigm as indicated in the Results and Figures. Baseline responses were recorded for 60 min before the delivery of the TBS paradigm. Experiments within an age were grouped depending on the success of inducing potentiation with a threshold set at 120% of the naïve fEPSP slope.

### Statistics

Statistical analyses were performed using Prism (GraphPad Software, Inc.; San Diego, CA). Data are presented throughout as the mean ± SEM. The minimal level of significance was set at p<0.05. The total number of animals and slices of each cohort (genotype and age) used in the 8T LTP experiments presented in Figures 1-8 are presented in Table 1. In other experiments, the number of slices is indicated in parenthesis in Figure 6, 9 and 10. The specific tests and the outcomes are indicated in the figure legends.

**Figure 1:**
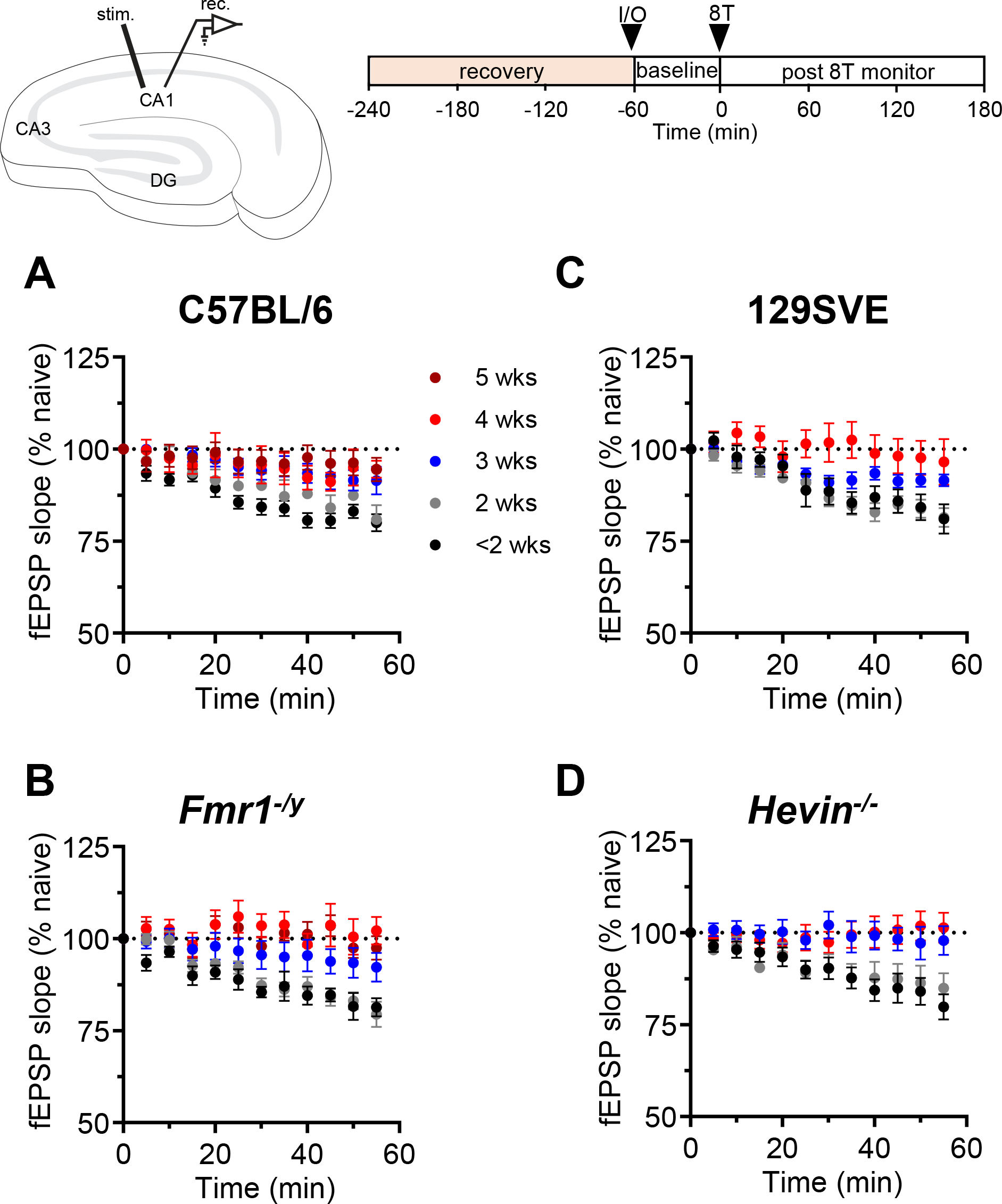
Test pulse-induced depression was resolved by 4-5 weeks for all strains and genotypes. (Top row) Electrode positions in hippocampal area CA1 and slice paradigm to test for L-LTP and experimental design. Slices were recovered for 3 hr without stimulation (tan frame). Then an I/O was done to determine the approximate half-maximal response (40-60%). The stimulus was repeated at this intensity at 5 min intervals for 60 minutes to obtain the baseline responses. The 8T consisted of 8 trains at 30 second intervals with 10 bursts at 5 Hz of 4 pulses each at 100 Hz. Then the responses were monitored for 180 minutes and L-LTP was determined by averaging the response slope over the last 155-180 min. All fEPSP slopes are normalized to the first (naïve) response at time 0. (A-D) baseline responses to the test pulse stimulation for the first 1 h of the experiments show an age-dependent decrease in test pulse-induced depression for all 4 genotypes (2-way ANOVA, Interaction: F(9, 208) = 0.427 (P=0.92), Age: F(3, 208) = 24.03 (P<0.0001), Genotype: F(3, 208) = 1.70 (P=0.17)). All genotypes revealed significant differences between ages before or at 2 weeks and 4-5 weeks old (Dunnet’s post-hoc tests). Data from 4 and 5-week-old mice are plotted together for *Hevin*^−/−^.

**Table 1:** Total number of slices and animals in each strain and age group for Figures 1-8. For Figures 9-10 the number of slices in each condition are indicated in parenthesis in the figures themselves.

| animals |  |  |  |  |  |
| --- | --- | --- | --- | --- | --- |
|  | <2wks | 2wks | 3wks | 4wks | 5wks |
| **C57BL/6** | 8 | 7 | 10 | 8 | 6 |
| ***Fmr1<sup>-y</sup>*** | 6 | 4 | 6 | 4 | 7 |
| **129SVE** | 6 | 6 | 6 | 6 |  |
| ***Hevin<sup>-/-</sup>*** | 5 | 5 | 6 | 7 | 4 |
| slices |  |  |  |  |  |
|  | <2wks | 2wks | 3wks | 4wks | 5wks |
| **C57BL/6** | 14 | 11 | 18 | 11 | 16 |
| ***Fmr1<sup>-y</sup>*** | 9 | 10 | 10 | 10 | 17 |
| **129SVE** | 10 | 11 | 10 | 10 |  |
| ***Hevin<sup>-/-</sup>*** | 11 | 12 | 13 | 14 | 10 |

## RESULTS

### Developmental test pulse depression

In immature rat hippocampus (<P20), test pulse stimulation produces a marked depression of the fEPSPs (Abrahamsson et al., 2007, 2008; Cao & Harris, 2012; Xiao, Wasling, Hanse, & Gustafsson, 2004). Both tetanic stimulation and TBS can reverse this test pulse-induced depression, but they produce no potentiation above the first naïve response at these young ages. To calculate the magnitude of test pulse depression and discern between the reversal of depression and LTP, all responses were normalized relative to the first naïve response. Here test pulses were given at the lowest frequency of 1 pulse per 5 min that was tested in rats (Fig 1). For mice aged ≤ 2 weeks, the test pulse depression was significant at 55 minutes (Table 2). By 4-5 weeks, the depression was gone, and such age-dependence was similar between all four mouse strains and genotypes. Thus, like in rats, test pulse depression in mice was age dependent.

**Table 2:** The magnitude of test pulse depression across ages and genotypes. The responses were normalized to the first naïve slope and the difference between the normalized average slopes at time 0 and 55 min is shown with means and 95% confidence interval (upper and lower limits).

|  | **C57BL/6** |  |  | ***Fmr1<sup>-y</sup>*** |  |  | **129SVE** |  |  | ***Hevin<sup>-/-</sup>*** |  |  |
| --- | --- | --- | --- | --- | --- | --- | --- | --- | --- | --- | --- | --- |
|  | Mean | Upper Limit | Lower Limit | Mean | Upper Limit | Lower Limit | Mean | Upper Limit | Lower Limit | Mean | Upper Limit | Lower Limit |
| <2wks | -0.19 | -0.15 | -0.23 | -0.17 | -0.12 | -0.23 | -0.16 | -0.09 | -0.24 | -0.17 | -0.10 | -0.24 |
| 2 wks | -0.16 | -0.08 | -0.23 | -0.18 | -0.12 | -0.24 | -0.17 | -0.12 | -0.21 | -0.14 | -0.05 | -0.23 |
| 3 wks | -0.08 | -0.01 | -0.15 | -0.07 | 0.02 | -0.15 | -0.09 | -0.06 | -0.11 | -0.02 | 0.06 | -0.10 |
| 4-5 wks | -0.05 | -0.01 | -0.10 | 0.00 | 0.05 | -0.04 | -0.03 | 0.08 | -0.14 | 0.05 | 0.12 | -0.03 |

### Strain- and genotype-specific differences in developmental onset of STP and L-LTP

In young adult rats and C57BL/6 mice (7-9 wks old), L-LTP is saturated by eight trains of TBS (8T) delivered in *s. radiatum* of hippocampal area CA1 (Cao & Harris, 2014). Saturating in this context means that a repeat episode of TBS given five minutes after the first episode produces no additional LTP. This protocol reliably produced L-LTP at P12 in rats (Cao & Harris, 2012). Hence, we used this 8T protocol to determine the onset age of L-LTP in acute slices from mouse hippocampal area CA1 (Fig. 1, top right). The LTP threshold was set at 120% of the naïve response.

For each age, the number of experiments was tabulated where TBS failed (none) or succeeded in producing short-term potentiation lasting 1 hr (STP) or L-LTP lasting at least 3 hr (LTP). Age groupings were based on the relative frequency of slices showing L-LTP (Fig. 2-6). The *Before Onset* groups comprised ages where L-LTP was induced in 25% or less of tested slices. The *Onset* groups had 26-50% success rate among tested slices. The *After Onset* groups had L-LTP in more than 50% of tested slices. The percentage was calculated by dividing the number of slices that showed only STP (gray sectors) or L-LTP (red sectors) by the total number of slices tested in the corresponding age group as shown in pie charts in Figs 2 and 3.

**Figure 2:**
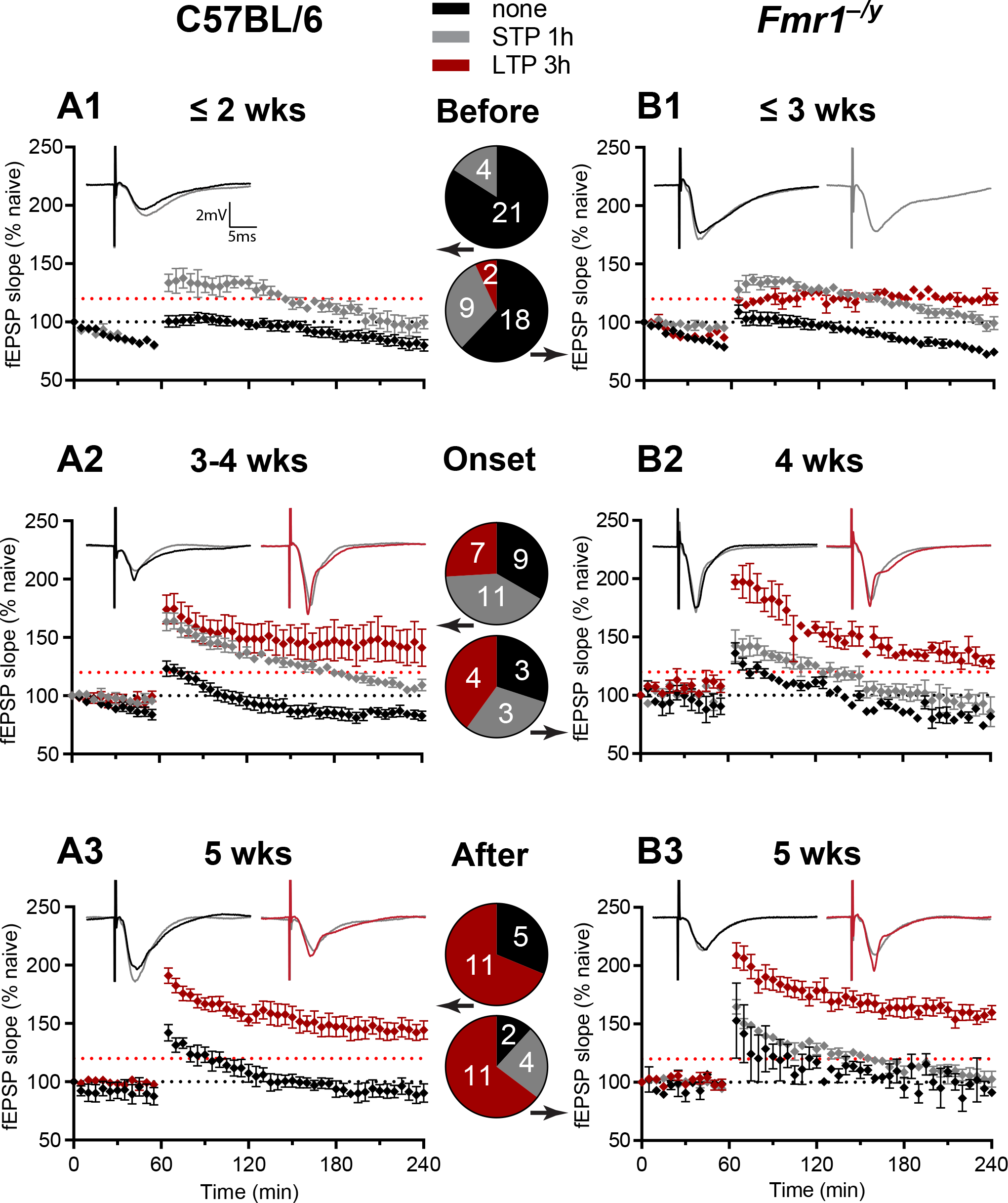
L-LTP was reliably produced by 5 weeks of age for (A) C57BL/6 and (B) *Fmr1*^*−/y*^ mice. (A1, B1) Before Onset age of L-LTP: Hippocampus from animals of both genotypes that were less than 3 weeks old rarely produced LTP lasting 3 hrs. (A2) Onset age of L-LTP: By 3-4 weeks, about a third of the C57BL/6 and (B2) nearly half of the *Fmr1*^*−/y*^ mice showed L-LTP lasting at least 3 hr. (A3, B3) After Onset age of L-LTP: By 5 weeks of age, well over half of the animals showed LTP lasting more than 3 hr for both genotypes. In time course plots and pie charts, the experiments with no potentiation (none) are colored black, those with STP lasting less than 1 hr (STP 1h) are colored gray, and those with LTP lasting 3 hr (LTP 3h) are colored red. Representative waveforms for pre-TBS baseline responses are colored gray, for 3 hr post-TBS are colored black for no potentiation and red for L-LTP at 3h. The pie charts show the relative fractions with the actual number of slices in each fraction for each age. The numbers of all animals and slices are also listed in Table 1.

**Figure 3:**
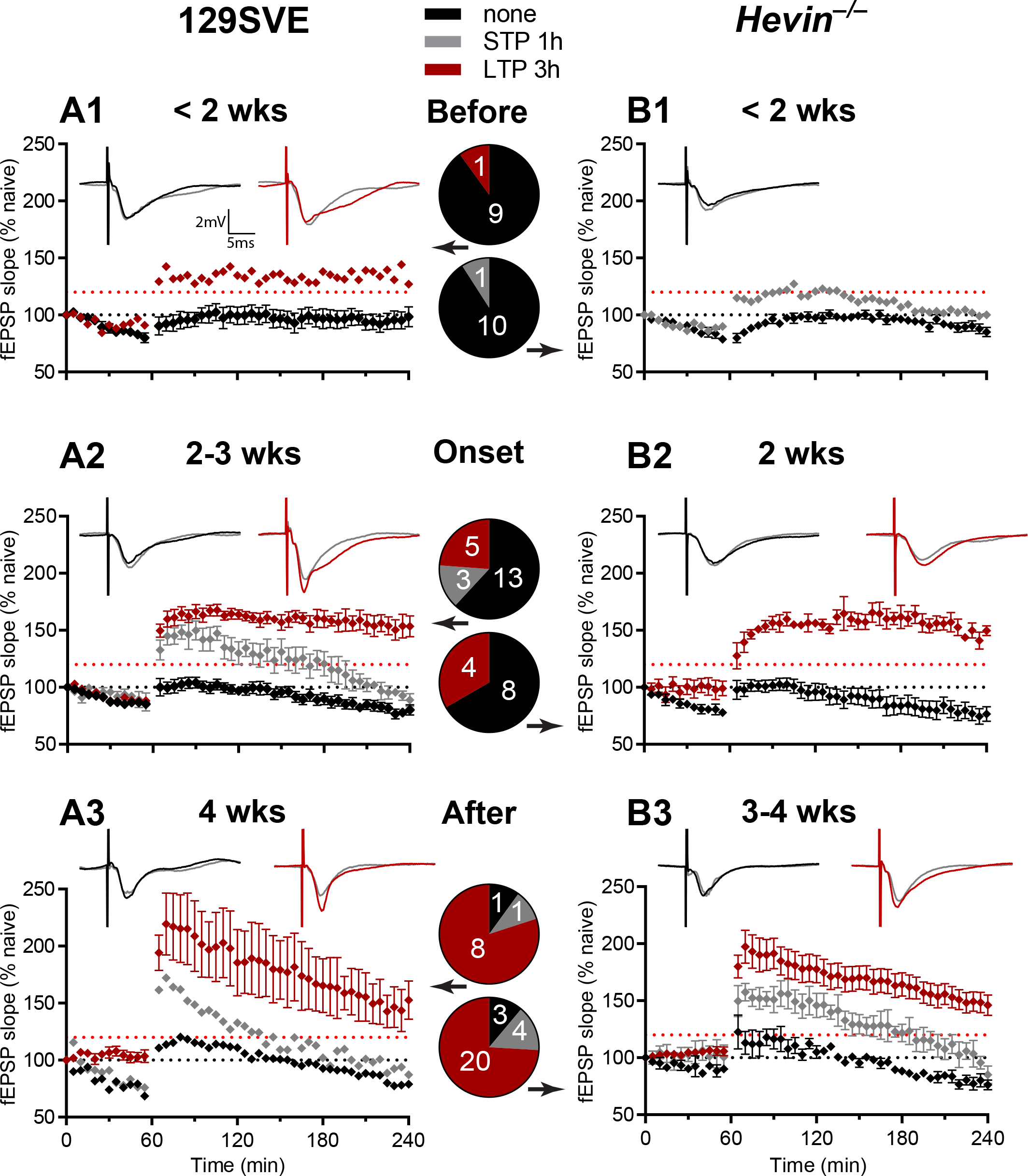
Week by week analysis of 1 hr and 3 hr LTP in the 129SVE (A1-4) and *Hevin*^−/−^ (B1-4) mice. The same color and labeling schemes as in Figure 1.

In mice from the C57BL/6 strain and *Fmr1*^*−/y*^ on C57BL/6 background, TBS reversed test pulse depression in all slices by 2 weeks before the onset of L-LTP (Fig. 2A1). By 2 weeks, about 16% of slices from the C57BL/6 wild type mice showed STP, but none had L-LTP (Fig. 2A1). By 3 weeks, 31% of slices from the *Fmr1*^*−/y*^ mutant mice showed only STP and 7% showed L-LTP (Fig. 2B1). Between 3-4 weeks, 41% of slices produced STP and 26% produced L-LTP in the C57BL/6 wild type (Fig. 2A2); whereas, in the *Fmr1*^*−/y*^ mutants, at 4 weeks 30% of slices produced STP and 40% produced L-LTP (Fig. 2B2). By 5 weeks, L-LTP was reliably produced in more than 50% of slices from both the C57BL/6 wild types and *Fmr1*^*−/y*^ mutants (Fig. 2A3, 2B3). Supplemental Figure 1 illustrates these findings on a weekly basis for each strain. These findings suggest a gradual onset for the production of STP and L-LTP in the C57BL/6 strain, an effect that was apparently delayed in *Fmr1*^*−/y*^.

Next, we tested mice from the 129SVE strain and *Hevin*^−/−^ on the 129SVE background. Before 2 weeks, all slices showed reversal of test pulse depression, one out of 10 slices showed minimal L-LTP in the 129SVE mice (Fig. 3A1), and one out of 11 slices showed subtle STP in *Hevin*^−/−^ (Fig. 3B1), but most slices showed no potentiation at all. By 3 weeks, 38% of slices showed either STP or L-LTP in the 129SVE mice (14% STP and 24% L-LTP, Fig. 3A2), contrasting with 33% of slices from *Hevin*^−/−^ showing L-LTP a week earlier by 2 weeks (Fig. 3B2). By 4 weeks, 80% of slices showed L-LTP and 10% had STP in the 129SVE mice vs 74% and 15% in *Hevin*^−/−^ (Fig. 3A3). Supplemental Figure 2 illustrates these findings on a weekly basis. Thus, reliable L-LTP occurred at 4 weeks in the 129SVE strain and even earlier, at 3 weeks, for *Hevin*^−/−^ (Fig. 3B3).

Additional experiments demonstrated that 1T, 2T, and 8T produced L-LTP of the same magnitude and endurance at 4-5 weeks (Fig. 4). Hence, 8T was a conservative saturating approach consistent with the same paradigm that saturated L-LTP in adult rats and adult C57BL/6 mice (Abraham & Huggett, 1997; Cao & Harris, 2014; Kramar et al., 2012).

**Figure 4:**
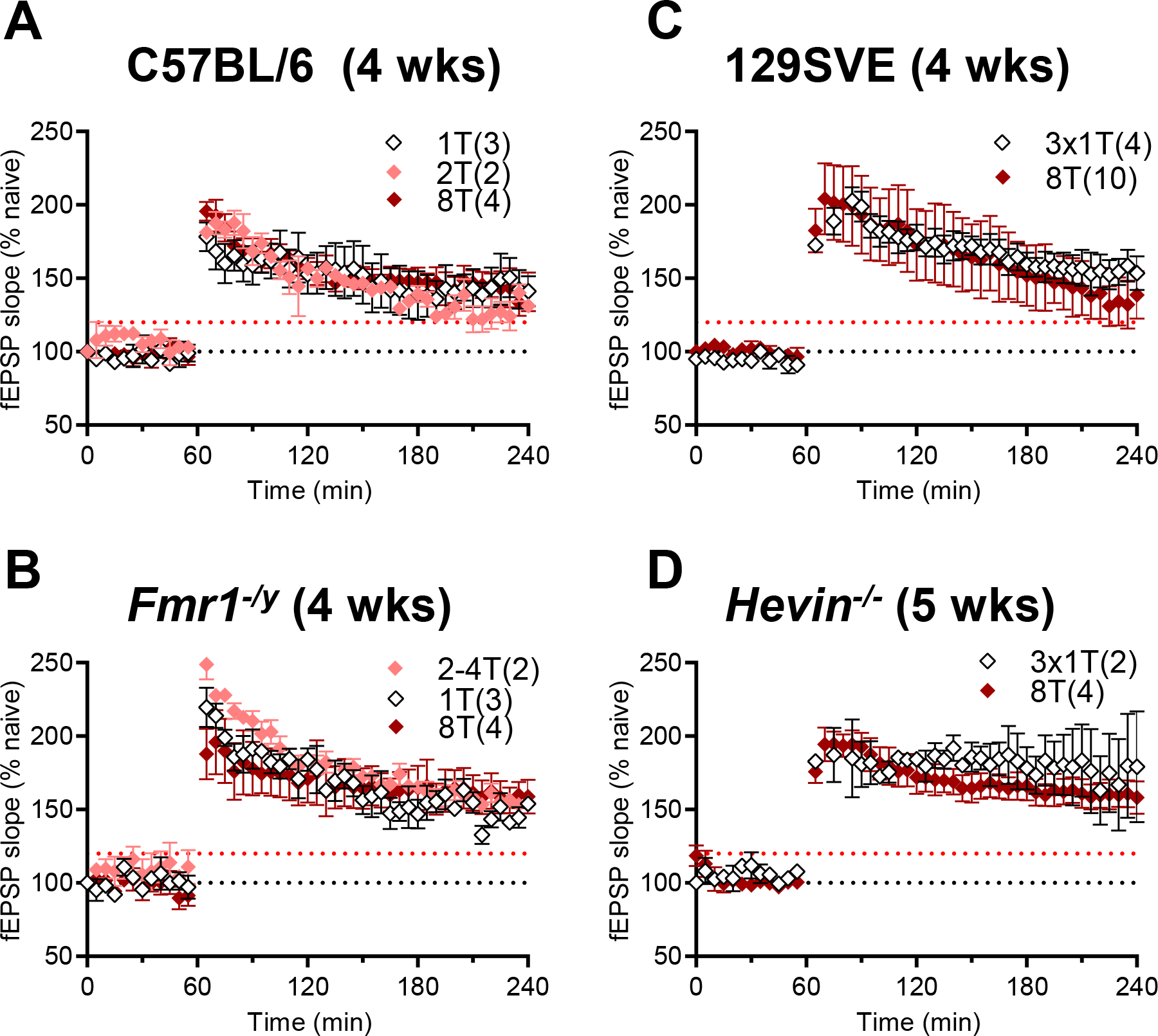
Demonstration that 8T is a robust induction paradigm for L-LTP for all strains and genotypes. Reducing the number of trains in the TBS paradigm from 8T to 1-4T resulted in the same magnitude and endurance of L-LTP in (A) C57BL/6, (B) *Fmr1*^*−/y*^, (C) 129SVE, and (D) *Hevin*^−/−^. The slices were obtained from animals at After Onset ages for each genotype.

### Both Fmr1^−/y^ and Hevin^−/−^ accelerated L-LTP onset relative to their wildtype backgrounds

The probability of producing at least STP (including STP and L-LTP slices) or L-LTP was compared across ages, strains, and genotypes (Fig. 5). More slices showed STP in *Hevin*^−/−^ mice by week 3 than other strains; however, this effect was not statistically significant, and all strains had ≥ 50% of slices showing STP at this age (Fig. 5A). The Onset age for L-LTP was significantly earlier at 3 weeks for the *Hevin*^−/−^ (corresponds to the LTP probability between ~50 and 75%) and at 4 weeks for *Fmr1*^*−/y*^ vs C57BL/6 (Fig. 5B). The After Onset age at 4 weeks in 129SVE and the *Hevin*^−/−^ (LTP probability ≥ 75%) was earlier when compared with 5 weeks in C57BL/6 and the *Fmr1*^*−/y*^ (Fig. 5B). The differences between C57BL/6 and *Fmr1*^*−/y*^ and between 129SVE and *Hevin*^−/−^ were also significant at 3 weeks (Fig 5B). Once established, there were no significant differences in the magnitude of L-LTP between mouse strains or genotypes across time post-induction (Fig. 5C).

**Figure 5:**
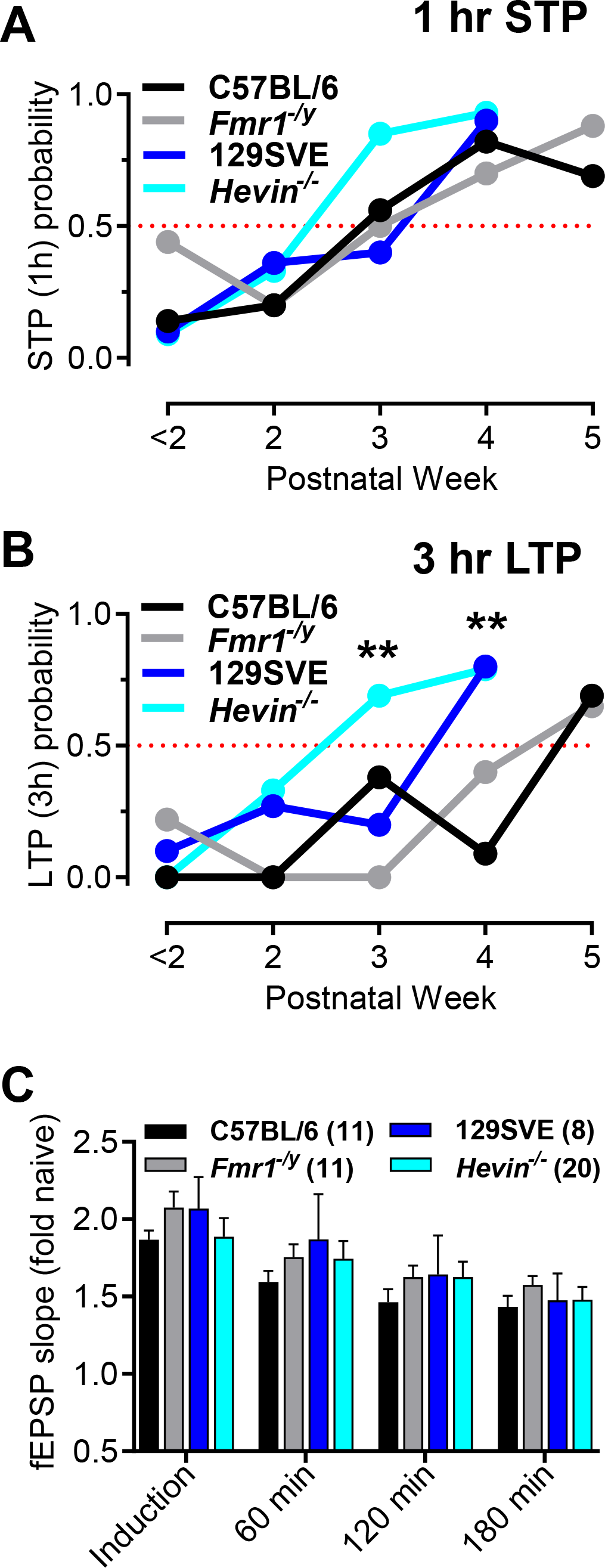
Differences among strains in the probability of L-LTP. The probability of STP (A) and L-LTP (B) by strain and genotype across postnatal age. The probabilities were calculated as the ratio of the number of successful STP or L-LTP experiments relative to the total number of experiments for each condition and age group. No significant differences were detected for STP. Significant differences were detected for the probability of L-LTP (B) at postnatal week 3 (χ^2^ = 13.16, df = 3; **P=0.0043) and postnatal week 4 (χ^2^ = 16, df = 3; **P=0.0012). 129SVE strain had a significantly higher probability of L-LTP than C57BL/6 at 4 wks (χ^2^ = 12, df = 3; ***P=0.0006). Within backgrounds, a significant difference was detected at 3 weeks between C57BL/6 and *Fmr1*^*−/y*^ (χ^2^ = 4.875, df = 1; *P=0.0272) and between 129SVE and *Hevin*^−/−^ (χ^2^ = 5.490, df = 1; *P=0.0191). All genotype pairs were equal at week 5. (C) The magnitude of fEPSP potentiation at different time periods post TBS did not differ significantly across strains or genotypes in the After Onset age groups for each strain or genotype.

Another measure of developmental onset would be the coincidence of no potentiation and potentiation occurring in different slices from the same animal. This coincidence was calculated as the percentage of animals that had slices showing both no potentiation and potentiation lasting for at least 1 hr out of the total number of animals at the Onset age for each genotype (Table 3). More than 60% of animals at the L-LTP onset age showed this coincidence in support of the hypothesis that these were indeed the relative onset ages for each genotype.

**Table 3:** The percent of animals at the onset age of L-LTP from each genotype that had some slices showing no potentiation and other slices from the same animal showing at least one hour of potentiation (coincidence).

| **genotype** | **coincidence** |
| --- | --- |
| **C57BL/6** | 66% |
| ***Fmr1<sup>-y</sup>*** | 75% |
| **129SVE** | 75% |
| ***Hevin<sup>-/-</sup>*** | 60% |

### The magnitude of the naïve fEPSP did not predict the developmental onset of L-LTP

The naïve fEPSP slopes were compared across experiments to test their potential effect in determining when L-LTP was first produced. In each of the three key age groups, the slices were divided as having no L-LTP at 3 hr or having L-LTP at 3 hr. No significant differences in the naïve slopes were detected across strains, genotypes, or age groups (Fig. 6). Thus, the magnitude of the naïve responses did not determine the occurrence of L-LTP in these experiments.

**Figure 6:**
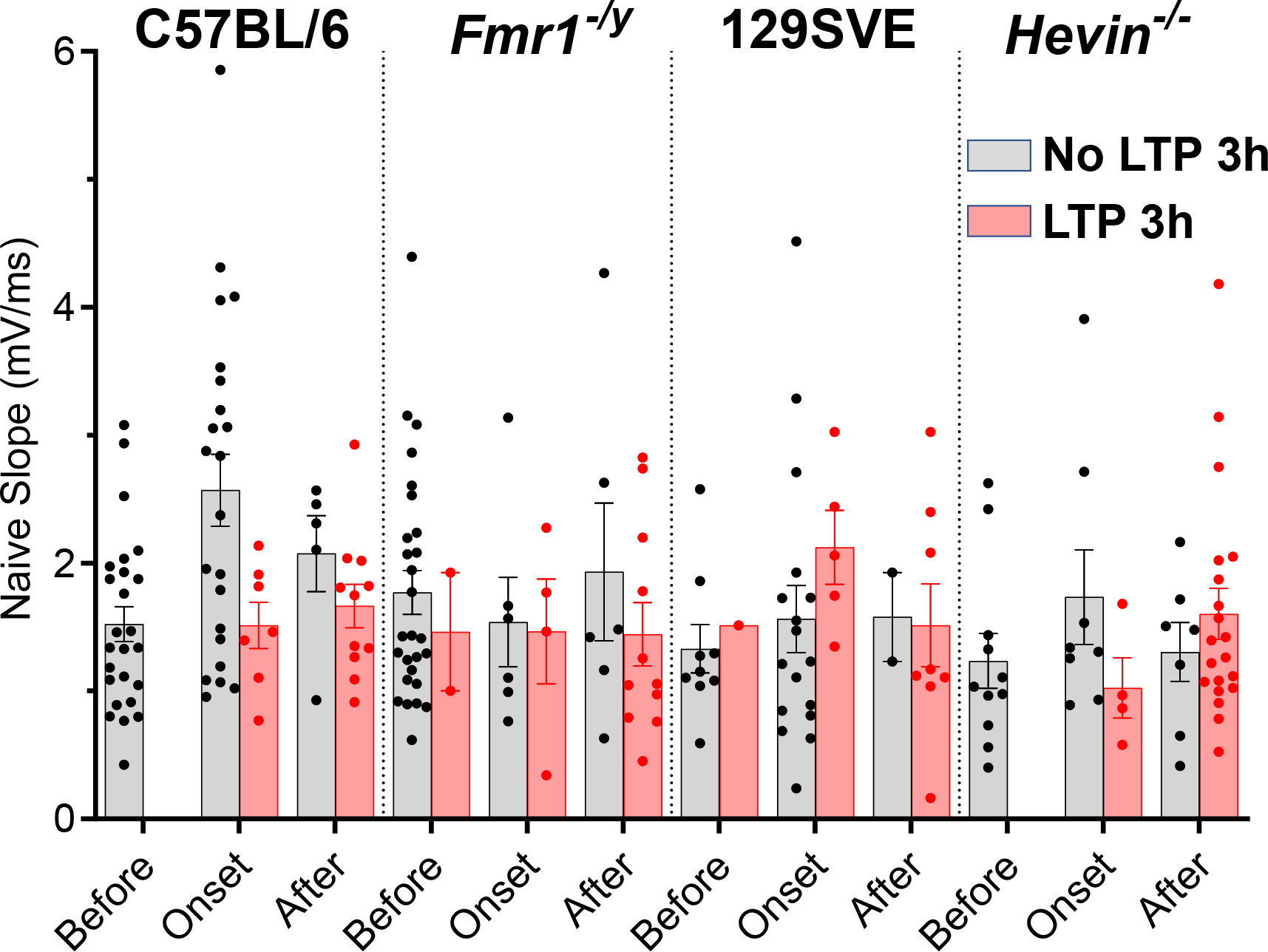
Magnitude of starting naïve responses did not predict success of L-LTP across ages, strains, or genotypes. Slices in each age group lacking L-LTP (black dots, gray bars) or producing 3 hr L-LTP (red dots and bars). Age groups are indicated as Before, Onset, and After the onset of L-LTP for each genotype. No significant differences were detected across the genotypes or stages for the Before Onset group, or for the Onset vs After L-LTP onset groups (2-way ANOVA, Interaction: F(7, 127) = 0.691 (P=0.680), Age: F(1, 127) = 0.0722 (P=0.789), Genotype: F(1, 127) = 1.57 (P=0.151)). Individual slice values are plotted as dots. Only 3 slices showed L-LTP in the Before Onset group, so these were not included in the statistical analyses.

### Age dependence of the augmentation of L-LTP

Prior work in the rat hippocampus revealed that multiple time-separated episodes of TBS result in additional potentiation or augmentation of L-LTP. The timing of this effect is strain and age-related in the rat hippocampus (Bowden, Abraham, & Harris, 2012; Cao & Harris, 2012, 2014; Kramar et al., 2012; Manahan-Vaughan, 2000; Manahan-Vaughan & Schwegler, 2011). We tested whether a pair of 8T episodes spaced 180 min apart would augment L-LTP in mouse hippocampus, as it does in rat (Fig. 7A). Two criteria were set for augmentation. First, the initial potentiation had to be ≥120% of the naïve response at three hours following the first 8T. Then, the second 8T had to elevate the fEPSP slope ≥10% above the initial L-LTP, and last for at least 70 minutes. Longer monitoring of the responses in slices from developing animals could become unreliable after 11 hours *in vitro* so experiments were terminated by 9 hours (i.e., 3 hours recovery plus 6 hours of recording).

**Figure 7:**
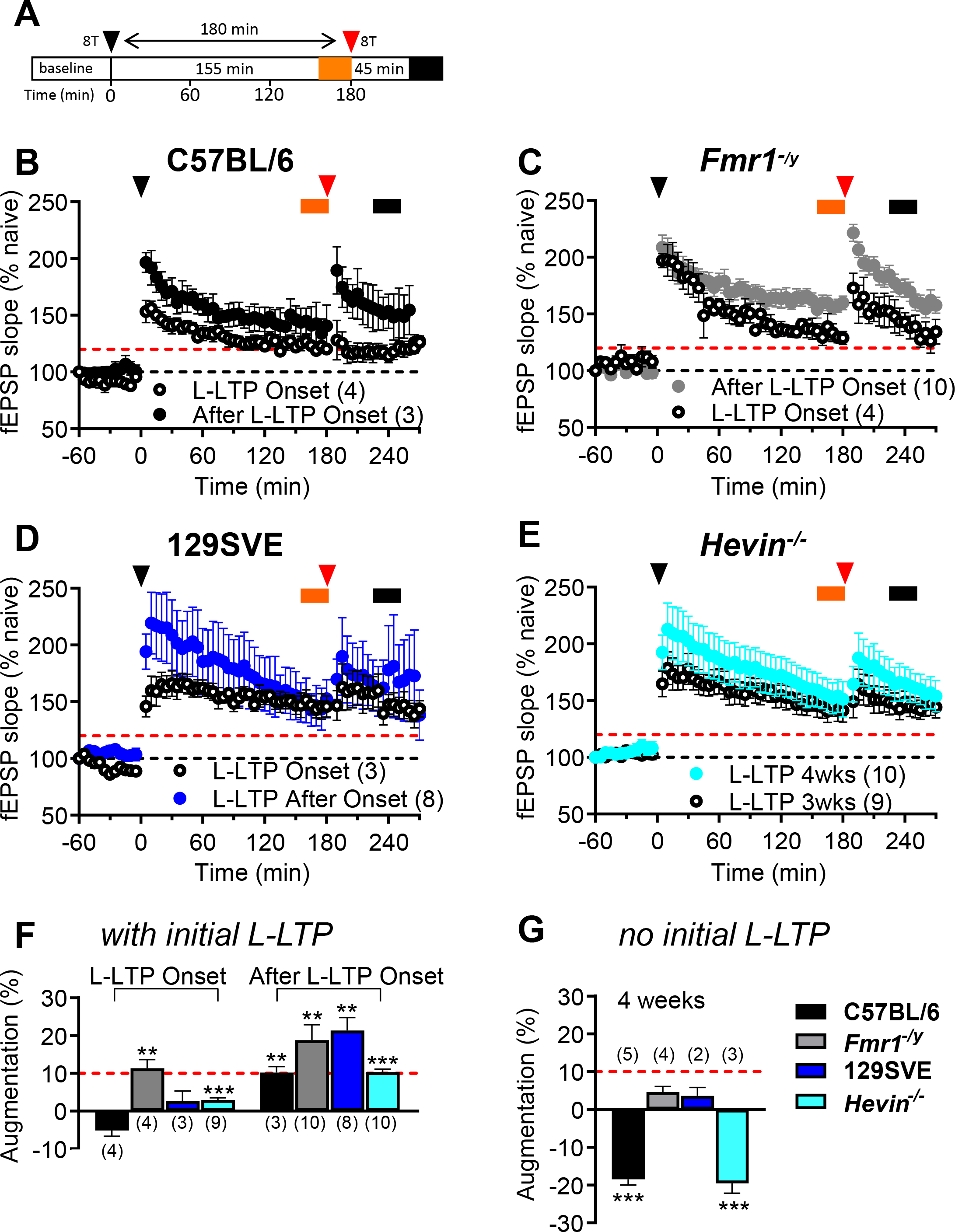
A second episode of 8T separated in time augments the initial L-LTP. (A) Experimental schematic: Recovery, I/O, Baseline, and first 8T matched original experiments (see Figure 1). Responses were monitored for 180 minutes after the first 8T, and a second 8T was delivered at 180 min (red arrowhead). The initial L-LTP was averaged over 155-180 min after the first 8T (orange frame). The effect of the second 8T was calculated 45-70 min later (black frame, 225-260 min after the first 8T). (B-E) Normalized fEPSP slope time course during the experiment plotted for (B) C57BL/6, (C) *Fmr1*^*−/y*^, (D) 129SVE, and (E) *Hevin*^−/−^. Augmentation of L-LTP was calculated as the percentage difference between fEPSP slope after the second TBS (black frame) relative to that after the first TBS (orange frame). L-LTP was considered to have undergone augmentation if the difference was at least 10% (red dotted line). (F) Summary for the augmentation of LTP across genotypes at L-LTP Onset (C57BL/6: t=6.335, df=5, partial η^2^ = 0.889, **P= 0.0014; *Fmr1*^*−/y*^: t=4.646, df=5, partial η^2^ = 0.812; *P= 0.0056; 129SVE: t=6.221, df=5, partial η^2^ = 0.886, **P= 0.0016; *Hevin*^−/−^: t=5.184, df=5, partial η^2^ = 0.843, **P= 0.0035) and After Onset developmental stages in slices that expressed L-LTP. For *Hevin*^−/−^ the Onset group includes slices from 3 week old animals, and the After L-LTP Onset group included slices from the 4 week old cohort. (G) Lack of augmentation in slices from 4 week old animals that did not express L-LTP after the first TBS, and two genotypes showed further significant decline (C57BL/6: t=12.11, df=5, partial η^2^ = 0.967, ***P< 0.0001; *Hevin*^−/−^ : t=7.369, df=5, partial η^2^ = 0.916, ***P= 0.0007). For the *Hevin*^−/−^, slices from both 3 and 4 week old mice were technically in the After L-LTP Onset age group. However, slices from 4 week old *Hevin*^−/−^ animals produced a more robust augmentation of L-LTP, while slices from 3 week old *Hevin*^−/−^ animals produced less augmentation that did not reach the criterion (Fig. 9E,F). Hence, the *Hevin*^−/−^ slices tested for augmentation were separated by age, with 3 weeks in the Onset group to illustrate the modest augmentation and 4 weeks in the After Onset group to illustrate the more robust augmentation.

Slices from the different mouse genotypes were compared in their respective Onset or After Onset of L-LTP age groups. The time course of L-LTP and augmentation of L-LTP is illustrated for all four genotypes (Fig. 7B-E). At the age of L-LTP Onset some slices from the *Fmr1*^*−/y*^ mutant mice, which had threshold levels of initial L-LTP, produced augmentation of L-LTP (Fig. 7C, F). None of the other genotypes met the 10% augmentation criterion (Fig. 7F). After the age of L-LTP Onset, all 4 genotypes produced augmentation of L-LTP (Fig. 7E-F). To illustrate the age-dependent difference in augmentation, the *Hevin*^−/−^ slices tested for augmentation were separated by age, with 3 weeks in the Onset group and 4 weeks in the After Onset group (Fig. 7F).

Even after the onset age of L-LTP, some slices showed no initial L-LTP. In those 4 week old slices, the second 8T also produced no potentiation and hence no augmentation (Fig. 7G). In fact, for slices without initial L-LTP in the C57BL/6 and *Hevin*^−/−^ mice, the second 8T episode resulted in a significant reduction in fEPSP slope 45-70 minutes later (Fig. 7G). This depression could merely reflect the failure to reverse the ongoing decline in the fEPSP slope normally observed over time in slices from developing animals that fail to produce initial L-LTP.

### Multiple episodes of 8T do not enable L-LTP in slices initially lacking production of L-LTP

Since two episodes of 8T produced no potentiation in slices lacking initial L-LTP, further testing was done varying the timing (90 vs 180 min intervals) and strength (8T vs 1T) of the episodes (Fig. 8A). Four-week-old C57BL/6 mice were chosen as this age was at the onset when a single episode of 8T did not reliably produce L-LTP. When two episodes of 8T were spaced by 90 minutes or 180 minutes, and STP was present initially for 1 hour,no additional STP was produced after the second 8T and the response dropped back to baseline or below by 3 hours (Fig. 8B,C). When the gentler 1T episodes were spaced by 90 minutes, substantial potentiation was induced after both the first and the second 1T episodes; however, it did not last (Fig. 8D, E). If the 1T episodes were spaced by 180 minutes, the second episode produced much less potentiation than the first, and it also did not last (Fig. 8D,E). Thus, neither change in timing or strength produced reliable initial L-LTP nor augmentation of STP to produce L-LTP at the earlier developmental stage.

**Figure 8:**
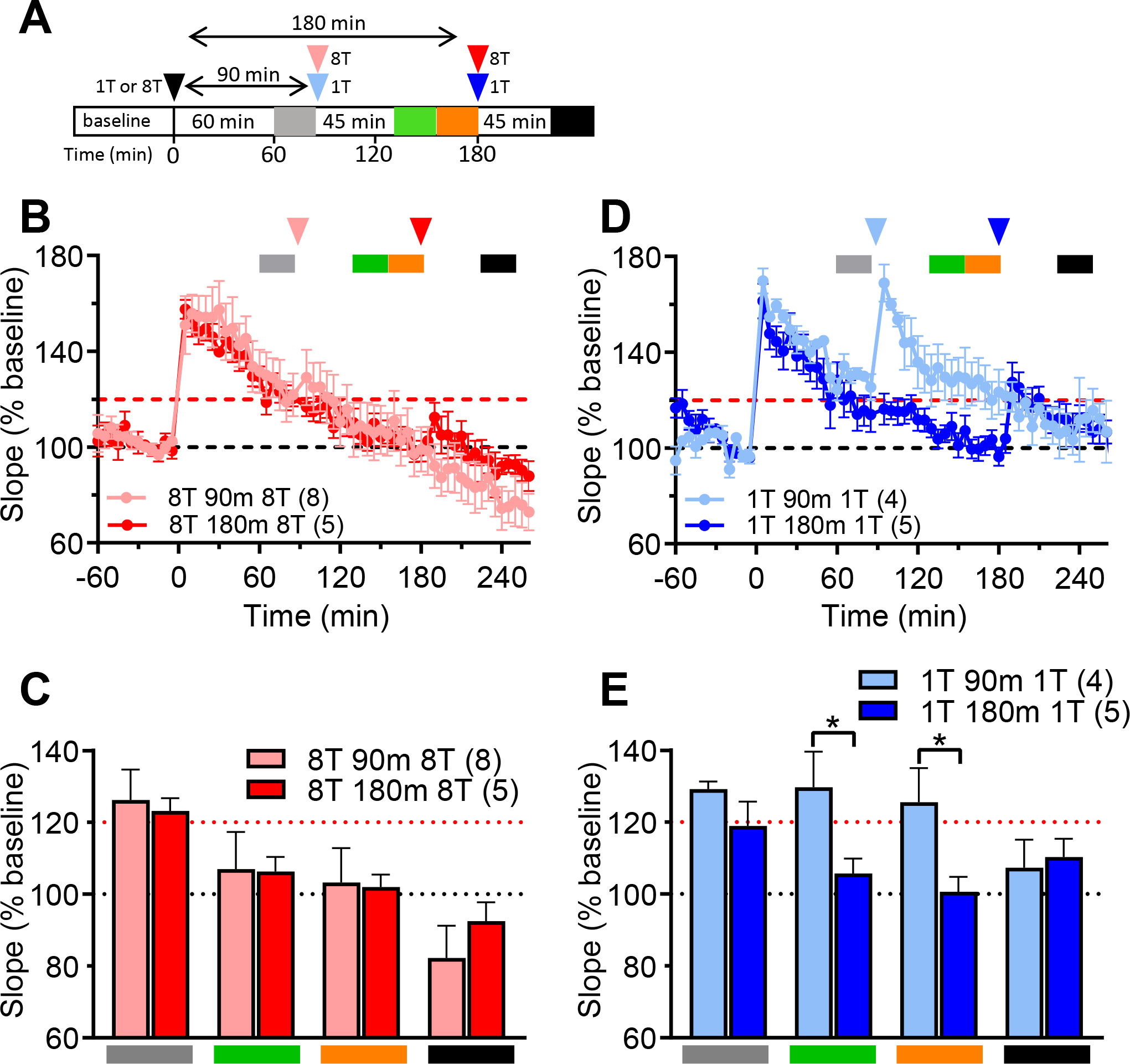
Three episodes of 8T did not produce L-LTP in slices lacking initial L-LTP in 4-week-old C57BL/6 mice. (A) Experimental schematic: Each slice was subjected to two identical TBS paradigms consisting of either 1 or 8 trains that were spaced by 90 min (1T light blue, 8T pink arrows) or 180 min (1T navy, 8T red arrows). Initial L-LTP was calculated by averaging responses 60-85 min before the second 8T at 85 min (gray time frame) or 155-180 min for the 180 min 8T interval (orange time frame). The effect of the second 8T was calculated at 135-160 min for the 90 min interval (green time frame) or at 225-260 min for the 180 min interval (black time frame). (B) Summary of the mean changes in fEPSP slope normalized to the 30 min averaged baseline responses with 8T TBS episodes spaced 90 min (pink) or 180 min (red). (C) Quantification of the experiments at the representative time frames for the experiments in B. No significant differences were detected at any of the time points for the different separations in 8T. (D) Summary of the mean changes in fEPSP slope normalized to the 30 min of the averaged baseline responses with 1T TBS episodes spaced 90 min (blue) or 180 min (navy). (E) Quantification of the experiments at the representative time frames for the experiments in D. No significant differences were detected between the levels of potentiation at orange and black intervals (pre and post second 1T spaced 180 min after the first 1T, 1-way ANOVA, F (3, 9) = 2.736, P = 0.267). A significant difference was detected by 2-way RM ANOVA between 1T-90m-1T and 1T-180m-8T by TBS spacing factor (F (1, 7) = 6.762; *p=0.0354).

## DISCUSSION

In the past, two major induction protocols have been used to discern the developmental onset of synaptic plasticity in the rat hippocampus. Repeated tetanic stimulation (3 times at 100 Hz for one second each) first produced L-LTP at P15 (K.M. Harris & Teyler, 1984; Jackson, Suppes, & Harris, 1993), while the 8T paradigm produced L-LTP earlier, at P12 (Cao & Harris, 2012). Hence, the more efficient 8T paradigm was adopted here to investigate the developmental onset of plasticity in the mouse hippocampus. In rats, STP and L-LTP developmental onset occurred at the same time. In contrast, three stages of plasticity emerged in mouse hippocampus. First, STP lasting only 1 hour gradually emerged between P10-P28 in all four mouse genotypes. Second, the Onset age of L-LTP (25-50% of slices) occurred at different ages depending on mouse genotype. Third, the After Onset age L-LTP (>50% of slices) also occurred at different ages depending on mouse genotype. In the 129SVE mice, L-LTP Onset occurred by 3 weeks, reliable L-LTP was achieved by 4 weeks, and *Hevin*^−/−^ advanced this profile by one week. In the C57BL/6 mice, L-LTP onset occurred significantly later between 3-4 weeks, and reliable L-LTP was not achieved until 5 weeks. Although some of the *Fmr1*^*−/y*^ mice showed L-LTP before 3 weeks, reliable L-LTP was also not achieved until 5 weeks in this genotype. In rat hippocampus, multiple episodes of 8T advanced the onset age of L-LTP from P12 to P10. In contrast, multiple bouts of 8T did not elevate STP to L-LTP in mouse hippocampus, nor did it advance the Onset age of L-LTP in any mouse genotype. However, if L-LTP was produced, additional spaced bouts of 8T could augment its magnitude in the hippocampal slices of those mouse genotypes that were checked. Test pulse depression, indicated by a gradual decline in fEPSP magnitude, has been ascribed to reversable silencing of AMPARs (Abrahamsson et al., 2007, 2008). In rats and all four mouse genotypes, full reversal of test pulse depression occurred at five weeks, coincident with the latest Onset age of L-LTP in mice, but well after L-LTP onset in rats. Thus, differences between species, strains, and genotypes could not be explained by the capacity to avoid test pulse depression. Similarly, the baseline slopes of the fEPSPs were comparable across the species, genotypes and ages. Hence, the differences in L-LTP onset ages were not explained by differences in the initial strength of activation reported here for mice, or previously for rats (Cao and Harris, 2012).

### Developmental metaplasticity

Some patterns of stimulation have no direct effect on synaptic strength but instead modulate the subsequent expression of plasticity, a phenomenon known as metaplasticity (Abraham & Bear, 1996; Abraham & Tate, 1997; Young & Nguyen, 2005). Spaced learning produces longer memories than massed learning, and the efficacy of memory is dependent on the interval between episodes of learning (Ebbinghaus, 1885; Fields, 2005). Similarly, spacing episodes of plasticity induction is considered to be a good model for understanding the cellular mechanisms of spaced learning (Kramar et al., 2012; Lynch & Gall, 2013; Lynch, Kramar, Babayan, Rumbaugh, & Gall, 2013). Regarding LTP, sufficient time must pass between the TBS episodes to augment LTP after a second episode of TBS. In adult rat, augmentation of previously saturated LTP was first observed at a 90 min spacing between bouts of 8T, and prolonging the time between TBS episodes increased the probability of augmentation (Cao & Harris, 2014). The delay between episodes of TBS reflects the time needed to enlarge the postsynaptic area after the initial induction of LTP in adult rats (Bell et al., 2014). In adults, this synaptic enlargement is also homeostatically balanced by stalled spine outgrowth that reflects temporal dynamics of resource reallocation to clusters of potentiated synapses (Bell et al., 2014; J. Bourne & Harris, 2007; Chirillo, Waters, Lindsey, Bourne, & Harris, 2019).

The patterns of augmentation of L-LTP are also developmentally regulated. In rats, a second episode of 8T delivered 90 min after the first episode produced L-LTP at P10-P11 but not at P8-P9. In the C57BL/6 mice, applying a second 8T episode 90 or 180 minutes after the first did not produce L-LTP even at 4 weeks of age, when STP could be produced in most slices. Instead, in mouse hippocampus, augmentation could be achieved only after L-LTP was reliably established for C57BL/6, 129SVE, and *Hevin*^−/−^ genotypes at After Onset age. Curiously, the augmentation of L-LTP was observed earlier in *Fmr1*^*−/y*^ mice than other genotypes, in slices that had initial L-LTP. These observations suggest that the development of L-LTP and metaplasticity involve processes that depend on species, strain, and genotype, but are independent from those processes that result in STP.

### Genetic Differences between Mice and Rats

Such striking differences between mice and rats in their developmental profiles of synaptic plasticity are consistent with genetic analysis. Almost half of the ~1K genes tested so far also show differential expression between mouse and rat hippocampal dendrites, with much less divergence in the other tissues (Francis et al., 2014). There are also large differences between rat and mouse adult hippocampal neurogenesis, a process that is especially important for learning and memory (Lazarov & Hollands, 2016; Snyder et al., 2009). Rats have more adult-born, death-resistant neurons, and these neurons mature faster in rats than in mice. The young neurons show a much higher contribution to fear learning tasks in rats than mice (Miller & Hen, 2015). These genetic and functional differences are consistent with rats having an earlier and more discrete onset age of L-LTP than mice.

### Contrasting Developmental Onset of L-LTP and Spinogenesis in Rat and Mouse Hippocampus

In this work, we chose gene manipulations that had been reported to alter dendritic spines and synaptic plasticity. *Fmr1*^*−/y*^ neurons have been characterized by an overproduction of underdeveloped spines that might not support the plasticity events (He & Portera-Cailliau, 2013). Treatment of neonatal *Fmr1*^*−/y*^ mice with the antibiotic minocycline resulted in better learning outcomes along with enhanced spine maturation (Bilousova et al., 2009). Furthermore, in adult *Fmr1*^*−/y*^ mice (3-5 months old), spaced trials rescued learning deficits (Seese, Wang, Yao, Lynch, & Gall, 2014). In the developing *Fmr1*^*−/y*^ mice, however, STP was not augmented to L-LTP upon spaced bouts of 8T in slices that had no initial L-LTP. Thus, future work will be needed to know whether those spaced learning effects resulted from the augmentation of L-LTP or other processes.

Hevin is required for the development of thalamocortical connectivity between P14-P25 mouse cortex (Risher et al., 2014). Moreover, when Hevin is absent, cortical dendritic spines show significant immaturity demonstrated by fewer but longer spines, and a distinct refinement problem. In the second week of development cortical spines often receive innervations from one cortical and one thalamic axon. By P25, these multiply innervated spines are refined to receive either a thalamocortical or intracortical synapse in wildtype (129SVE) mice. In the *Hevin*^−/−^ mice this pruning effect does not occur uniformly and the ratio between thalamocortical and intracortical inputs is altered, retaining more of the intracortical synapses at the expense of thalamocortical connections. The role of Hevin in refining hippocampal dendritic spines is unknown; however, its absence in *Hevin*^−/−^ mice appears to advance the developmental onset age of L-LTP. This finding is consistent with the hypothesis that lack of refinement of CA3-CA1 synapses by Hevin promotes the earlier maturation of plasticity. Application of Hevin protein to autaptic cortical neurons results in a robust induction of NR2B containing NMDAR activity (Singh et al., 2016). Perhaps the lack of regulation of the NR2B subunit in the *Hevin*^−/−^ mice stimulates the maturation of synapses at an earlier developmental stage.

In rats, the developmental onset of L-LTP at P12 is coincident with the emergence of dendritic spines, suggesting dendritic spines are necessary for sustained synaptic plasticity (Cao & Harris, 2012; Fiala, Feinberg, Popov, & Harris, 1998; Kirov, Goddard, & Harris, 2004). Initial 3D reconstructions in perfusion-fixed rat hippocampus show evidence for mature dendritic spines at P12, but not at P8 or P10 (Smith, 2019). Preliminary data from rat hippocampal slices showed that 90 minutes after the initial 8T, dendritic spines were not produced at P8 (K. M. Harris, Watson, Kuwajima, & Cao, 2012). In rat hippocampal slices at P10-11, preliminary data suggest that spines were produced 90 minutes after the initial 8T (Smith, 2019). Thus, a shift from shaft synapses and filopodia to spines might account for this developmental shift in L-LTP onset for rat hippocampus.

The Onset of L-LTP in *Fmr1*^*−/y*^ at 4 weeks was delayed relative to the background C57BL/6 strain at 3 weeks. This pattern contrasted with the earlier onset age in *Hevin*^−/−^ by 2 weeks, relative to its background 129SVE strain between 2-3 weeks. All genotypes showed reliable L-LTP by 5 weeks. These findings contrast with the developmental onset ages of dendritic spines. Mature dendritic spines have been reported by P15 in C57BL/6 mouse hippocampus (Bilousova et al., 2009), well before the onset age of reliable L-LTP at 5 weeks. Confocal microscopy studies reveal a few mushroom spines by 9-12 days in organotypic slices from mouse hippocampus (Parnass, Tashiro, & Yuste, 2000). Similarly, at 14 days in organotypic slices from C57BL/6 and *Fmr1*^*−/y*^ more than 40% of the protrusions were classified as mushroom spines, although less than 10% had mature heads with a diameter greater than 0.5 μm (Bilousova et al., 2009). Reconstructions from serial section EM show mature spines by P24 in the C57BL/6 hippocampus (Nikonenko et al., 2013). Thus, the onset of L-LTP appears to be later than the onset of dendritic spines in mouse hippocampus, suggesting that spines might be necessary but not sufficient.

### Other factors that may influence variation in the developmental onset of L-LTP

The wide variance in L-LTP onset ages among individual mice may reflect divergence in many factors that lead to the maturation of neurons. In rats, the discrete onset age of L-LTP may reflect less variation in these factors between animals. Prior work also showed a high rate of failure of LTP induction in P16-P30 C57BL/6 mice when 5 tetanic stimuli were used in normal calcium concentration (Adesnik & Nicoll, 2007). The success rate of LTP induction was improved by increasing the calcium concentration. One possible explanation is the involvement of different mechanisms of LTP over developmental stages. There is evidence that in 2 week old mice of mixed 129SVE-C57BL/6 background the initial LTP is mediated by postsynaptic insertion of GluR2-lacking subunits which are later exchanged for GluR2-containing AMPA receptors that are less permeable to calcium (Jia et al., 1996; Plant et al., 2006; Purkey et al., 2018; Sanderson, Gorski, & Dell’Acqua, 2016). LTP in young mice is less dependent on the phosphorylation of GluR1 under mild stimulation conditions (Adesnik & Nicoll, 2007; Lee et al., 2003; Lu et al., 2007; Wikstrom, Matthews, Roberts, Collingridge, & Bortolotto, 2003). Thus, in addition to age, the exact induction protocol may influence the success of LTP.

Another factor that could influence the exact onset age of LTP is the maturation of the inhibitory system and the resulting excitation/inhibition (E/I) balance (Eichler & Meier, 2008) (Ben-Ari, Khalilov, Kahle, & Cherubini, 2012). The GABAergic system is depolarizing early during development but switches with maturation to hyperpolarizing. Varying the inter-stimulus interval shows that paired-pulse inhibition first occurs in the developing rat hippocampus at P6 (K.M. Harris & Teyler, 1983). Patch-clamp experiments, in hippocampus from both mice and rats, reveal that prior to the maturation of the GABAergic system, the degree of postsynaptic depolarization needed to induce LTP is less (Meredith, Floyer-Lea, & Paulsen, 2003). Furthermore, the *Fmr1*^*−/y*^ mice show a dramatic E/I imbalance mediated by reduced GABAergic inhibition in both hippocampus and subiculum (Cea-Del Rio & Huntsman, 2014; Curia, Papouin, Seguela, & Avoli, 2009; Eichler & Meier, 2008; Paluszkiewicz, Martin, & Huntsman, 2011; Sabanov et al., 2017). The saturating TBS protocol used here would overcome inhibitory effects at all ages; hence, the potential E/I imbalance would not explain the gradual onset of enduring LTP in mice (Pike, Meredith, Olding, & Paulsen, 1999; Thomas, Watabe, Moody, Makhinson, & O’Dell, 1998).

The gradual onset of L-LTP in mice could also stem from variation in the rate of maturation of neurons along the septal temporal axis (Altman, 1966; Altman & Das, 1966; Angevine, 1965; S.A. Bayer, 1980a, 1980b; S. A. Bayer & Altman, 1975). Taking four slices from the middle of the mouse hippocampus might overlap this developmental axis, whereas the larger rat hippocampus does not. This hypothesis is further supported by our finding that at onset, slices from the same mouse hippocampus could express no potentiation or potentiation lasting ≥1 hour. This coincidence occurred in a majority of mice from all genotypes at the onset ages of L-LTP. Such a finding might also reflect coincidental natural variation between neighboring hippocampal slices in the capacity for L-LTP and spine maturity. Thus, future experiments will need to measure both the capacity for L-LTP and spine structure in individual mouse slices to address the necessity and sufficiency of dendritic spines for L-LTP and the impact of these mutations on that process.

## Acknowledgements

We thank Dr. Darrin Brager for providing the breeding pairs of *Fmr1*^*−/y*^ mice, and Mr. Clayton Smith and Ms. Amanda Heatherly for maintaining the mouse colony.

