## Supplemental Figure 1 and 2 for "Developmental onset of enduring long-term potentiation in mouse hippocampus"

### Supplemental Fig 1

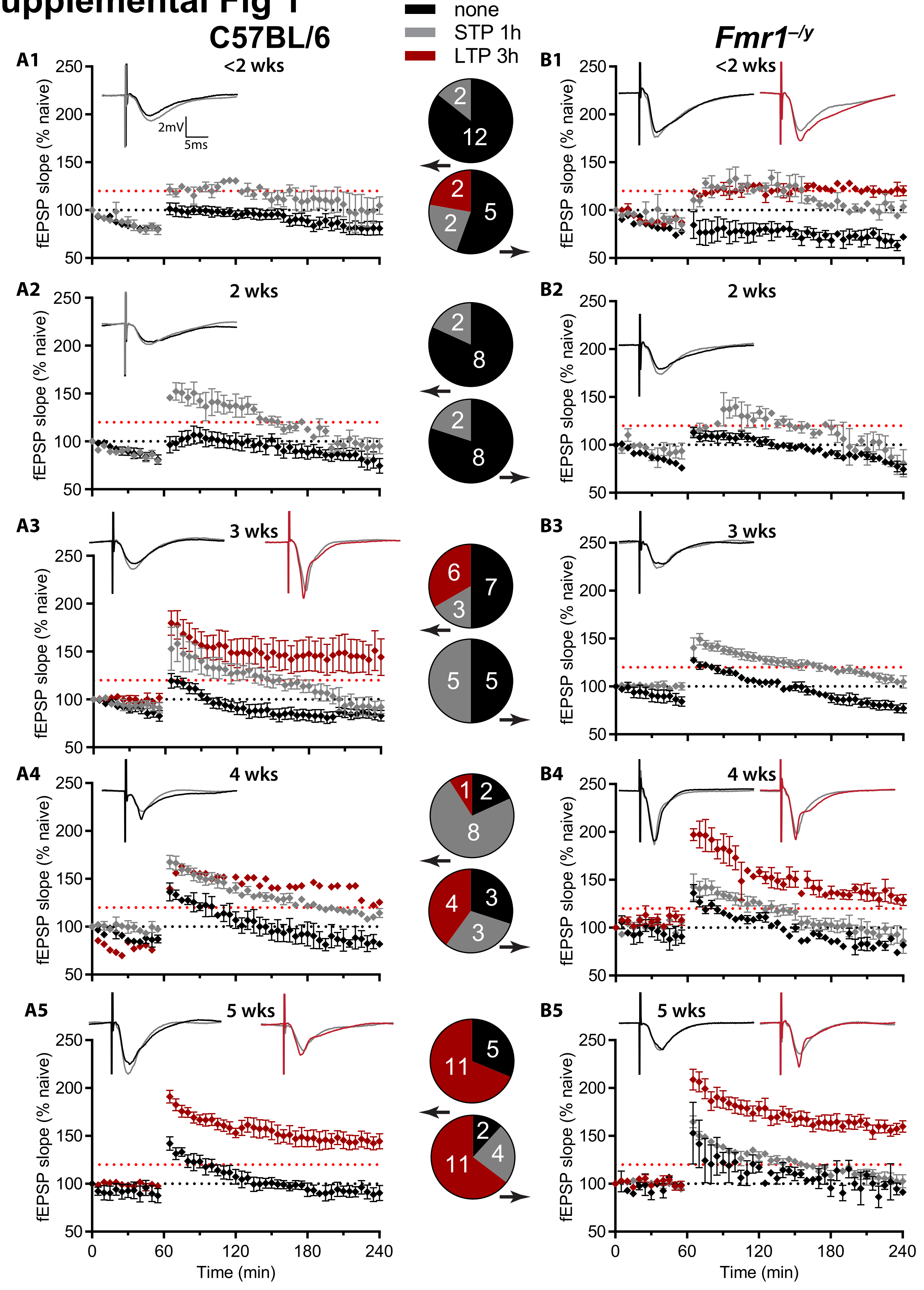

### Supplemental Fig 2

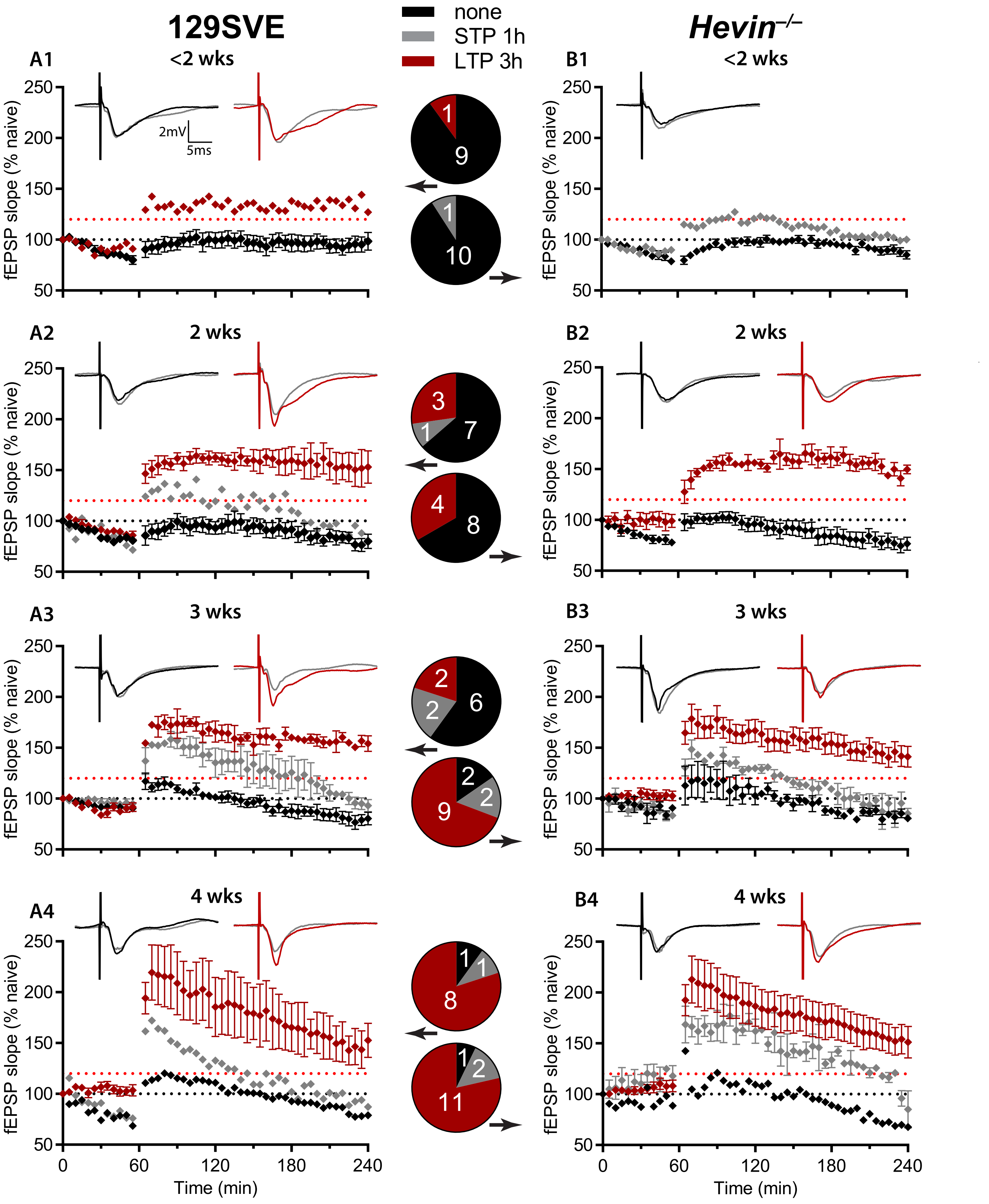

### Supplemental Figure Legends

**Supplemental Figure 1**: **Week by week analysis of STP and L-LTP in the C57BL/6 (A1-A5) and *Fmr1*$^{-/y}$ (B1-B5) mice.** The same color and labeling schemes as in Figure 1.

**Supplemental Figure 2: Week by week analysis of 1 hr and 3 hr LTP in the 129SVE (A1-4) and *Hevin*$^{-/-}$ (B1-4) mice.** The same color and labeling schemes as in Figure 1.
